# Heterogeneous associations between sex ratio distorters and mitochondrial haplotypes in U.S. populations of *Armadillidium vulgare*

**DOI:** 10.64898/2026.04.29.721737

**Authors:** Arushi Kansal, Robert Kuhn

**Affiliations:** FCS Innovation Academy STEM High School, Alpharetta, GA

## Abstract

Sex ratio distorters (SRDs) are heritable elements that bias offspring sex ratios to enhance their transmission. In the terrestrial isopod *Armadillidium vulgare*, feminization of genetic males can occur through vertical transmission of the sex ratio distorter known as the f-element, as well as through infection by *Wolbachia*, a maternally inherited bacterial endosymbiont that can alter host reproduction. Previous studies have focused on the distribution of SRDs and their associations with mitochondrial haplotypes in native European populations, but these patterns are poorly understood in the United States. In this study, we sampled *A. vulgare* in 12 states, screening individuals for *Wolbachia* infection, the presence of the f-element, and mitochondrial haplotypes. We found that *Wolbachia* shows a heterogeneous distribution across populations and haplotypes, in contrast with stronger associations in Europe. The f-element occurred in lower overall frequencies but showed a strong association with mitochondrial haplotype VI. These results indicate that patterns associated with SRD differ from those observed in Europe and suggest that multiple introductions and population mixing have shaped these distributions.

## Introduction

Sex ratio distorters (SRDs) are genetic elements or symbiotic microorganisms that alter host reproduction to bias offspring sex ratios, often increasing their transmission through host populations [1,2]. Many SRDs are passed maternally. One of the best-known examples is the endosymbiotic bacterium *Wolbachia pipientis*, which infects an estimated 50% of all arthropods [3]. *Wolbachia* manipulates host reproduction through four primary mechanisms. The most common is cytoplasmic incompatibility, in which male offspring from *Wolbachia*-infected males and uninfected females fail to develop. Male killing results in male offspring being killed by *Wolbachia*, resulting in only female offspring surviving. Parthenogenesis results in the production of females from unfertilized eggs, eliminating the need for males in reproduction. Feminization occurs when *Wolbachia* interferes with the development of the androgenic gland in genetic males, causing them to develop as reproductive females [2,3]

Sex determination in some terrestrial isopod species is variable and influenced by interactions with symbiont-derived elements [4], making *Armadillidium vulgare* a well-established system for studying *Wolbachia*-induced feminization. *A. vulgare* populations also include a nuclear sex-ratio distorter known as the f-element, which originated from the integration of a portion of the feminizing *Wolbachia* DNA into the host genome [5,6]. Because *Wolbachia* is maternally inherited while the f-element is integrated into the host nuclear genome, the two feminizing systems follow different inheritance patterns and evolutionary dynamics in natural populations.

*A. vulgare* is native to Europe but has been distributed to the U.S. through repeated human dispersal events associated with trade, agriculture, and urbanization [7]. In North America, population genetic studies indicate that *A. vulgare* did not arise from a single colonization event but instead reflects multiple independent introductions from genetically distinct European source populations. Mitochondrial DNA analyses show the coexistence of divergent COI lineages in U.S. populations [8]. Because these introductions occurred at different times across source populations, U.S. populations may preserve distinct mitochondrial haplotypes and varying frequencies of sex‐ratio distorters such as *Wolbachia* and the f-element. European surveys have shown clear associations between *Wolbachia* strains and COI haplotypes, as well as geographic structuring of SRD frequencies [9].

SRDs strongly influence population sex ratios and mitochondrial diversity in *A. vulgare*, yet their distribution in U.S. populations remains poorly characterized. European populations show heterogeneity of SRDs and non-random associations between feminizing elements and mitochondrial COI haplotypes, indicating strong host–symbiont dynamics [9]. In contrast, North American populations have not been surveyed at comparable geographic or genetic resolution. Preliminary laboratory work confirmed *Wolbachia*-associated male feminization in U.S. *A. vulgare*, demonstrating that feminizing symbionts actively distort sex determination in introduced populations [10], but the geographic extent of SRDs and their relationship to mitochondrial lineages in the wild is unknown. To address this gap, we conducted a nationwide sampling of U.S. *A. vulgare* populations, screened individuals for *Wolbachia* and the f-element, characterized mitochondrial COI haplotypes, and tested for associations between SRD presence and host mitochondrial haplotypes, thereby allowing comparisons with established European patterns.

## Materials and Methods

### Sample Collection

Pillbugs were obtained through direct collection by the authors and through citizen science crowdsourcing. We collected specimens from Arizona, Georgia, Hawaii, and Wisconsin. Specimens from California, Connecticut, Missouri, Nebraska, New York, North Carolina, Pennsylvania, and Washington state were obtained through crowdsourcing efforts coordinated via social media platforms. Participants collected and shipped specimens preserved in alcohol to our laboratory (Table 1). Once received, all pillbugs were preserved in 90% ethanol and stored in our lab at −20 °C. Specimens were visually identified as *A. vulgare* based on external anatomy and sexed by examining the ventral side for the presence or absence of the male endopods and later confirmed through COI barcoding using standard primers (LCO1490/HCO2198), and sequences compared against reference databases for species identification.

**Table 1.**
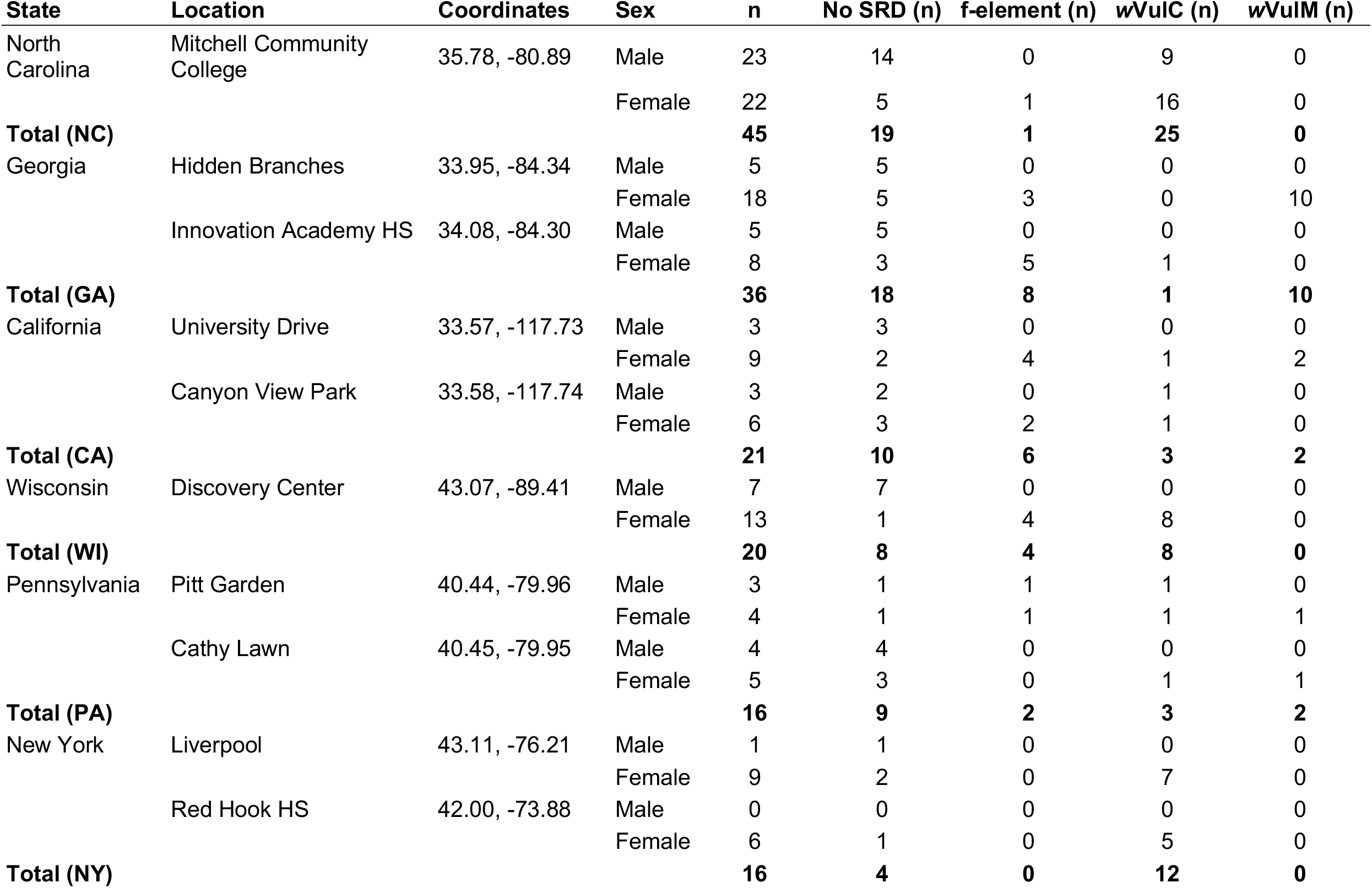
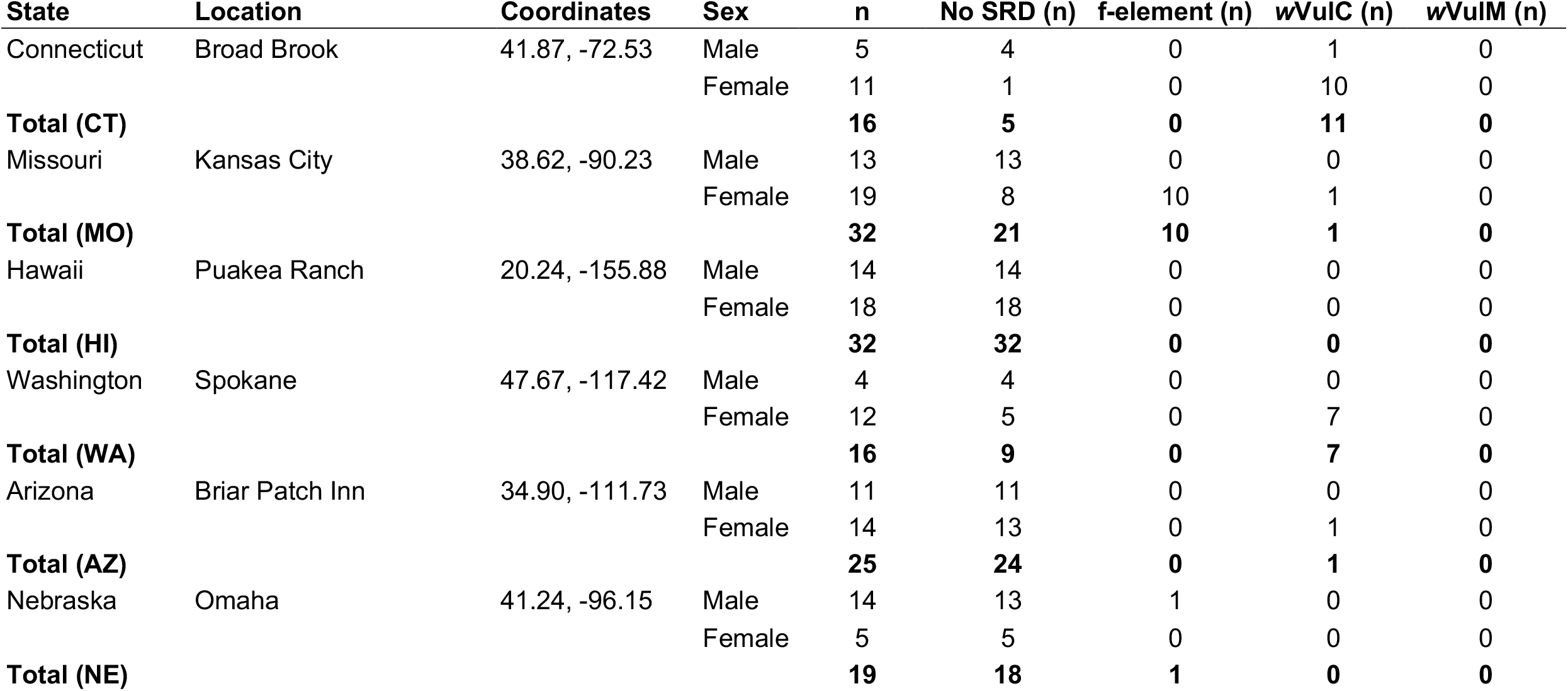
Distribution of *Wolbachia* and the f-element across sampling locations in the United States.

### DNA Extraction

Pillbug heads and legs were dissected for DNA extraction. Because *Wolbachia* is distributed systemically in *A. vulgare*, including in nervous tissue and hemolymph, it can be detected in non-reproductive tissues [11]. Head tissue was preferred, with leg tissue serving as an alternative when heads were not available. Both tissue types also yielded sufficient DNA for amplification of COI and detection of the f-element.

From 2023 to 2024, genomic DNA was extracted and purified using the ZymoBIOMICS DNA Miniprep Kit (D4300). Dissected tissues were first crushed in lysis buffer using a sterile micropestle, then transferred to bead bashing tubes containing a mixture of 0.5 and 1.0 mm silicon beads (as provided in the kit). Samples were homogenized by vortexing at maximum speed for 20 minutes using a Vortex-Genie 2. Following homogenization, samples were centrifuged (12,000 × g, 1 min), and the supernatant was used for downstream spin column DNA purification and elution according to the manufacturer’s protocol.

Beginning in 2024, genomic DNA extractions were performed using the Qiagen DNeasy Blood and Tissue Kit (69504) following the manufacturer’s *Purification of Total DNA from Insects* protocol with modifications. Dissected tissues were mechanically homogenized using a sterile plastic micropestle in buffer ATL rather than PBS, then centrifuged (12,000 × g, 1 min). The supernatant was transferred to a clean microcentrifuge tube, after which buffer AL and Proteinase K were added, and samples were incubated overnight at 57 °C in a shaking heat block. The resulting lysate was then processed according to the manufacturer’s protocol for spin column DNA purification and elution.

### Polymerase Chain Reaction (PCR)

To verify pillbug taxonomy and compare mitochondrial haplotypes, the mitochondrial DNA cytochrome c oxidase subunit I (COI) region was amplified. The COI locus was selected due to its widespread use as a species-level DNA barcode [12]. The primers used were LCO1490 (5′-GTAAAACGACGGCCAGGGTCAACAAATCATAAAGATATTGG-3′) and HCO2198 (5′-CAGGAAACAGCTATGACTAAACTTCAGGGTGACCAAAAAATCA-3′), producing an expected amplicon size of 700 bp.

Detection of *Wolbachia* infection was performed by amplification of the *Wolbachia* surface protein (*WSP)* gene using primers 81F (5′-TGGTCCAATAAGTGATGAAGAAAC-3′) and 691R (5′-AAAAATTAAACGCTACTCCA-3′), generating an expected 650 bp product [13].

The presence of the f-element nuclear insert was assessed using primers SubF1 (5′-ACGAAAAGCCGACGTAAAATATTT-3′) and JTelR2 (5′-GAAATAAAAGAGCCTGACT-3′), yielding an expected amplicon of approximately 700 bp.

PCR reactions were performed in a total volume of 35μL using Promega GoTaq 2× Green Master Mix (M3001), Promega nuclease-free water (MC1191), primers (10 μM), and template DNA. Each reaction contained 17.5 μL master mix, 11.9 μL water, 1.4 μL forward and reverse primers, and 2 μL of DNA.

WSP and f-element PCR conditions consisted of initial denaturing at 94 °C for 3 min, followed by 35 cycles of 94 °C for 30 s denaturing, 55 °C for 30 s annealing, 72 °C for 1 min elongation, and 72 °C for 10 min final extension [5]. COI PCR temperatures consisted of initial denaturing at 94 °C for 2 min, 30 cycles of 94 °C for 30 s, 49 °C for 45 s annealing, 72 °C for 1 min extension, with a final extension at 72 °C for 10 min.

### Sequencing and Bioinformatics

Visualization of PCR products was performed using 1.5% agarose gels, stained with GelGreen DNA stain (MiniPCR) under blue light illumination to confirm the expected amplicon size. Unpurified PCR products were submitted for Sanger sequencing, with cleanup performed by Azenta Life Sciences (GeneWiz) (South Plainfield, NJ, U.S.A).

Sanger sequences were analyzed using Geneious Prime (v2025.2). *A. vulgare* COI haplotypes were assigned by aligning Sanger sequences to 23 mitochondrial reference haplotypes [9], with each sample assigned to the closest matching reference based on highest sequence similarity. Overall, sequence identity was high, with 240 samples showing 99.0–100.0% identity, including 159 exact matches. An additional 36 samples showed 98.0–99.0% identity, 12 showed 97.0– 98.0% identity, and 5 showed 96.0–97.0% identity. Lower-identity sequences remained consistent with haplotype assignments, as they matched the same reference haplotypes observed in other individuals from the same populations with higher sequence identity.

*Wolbachia* strain identity was determined using NCBI BLAST searches of Sanger sequences, with assignments based on the highest sequence similarity to reference sequences. Sequence identity was high overall, with 69 samples showing 99.0–100.0% identity, including 25 exact matches. An additional 12 samples showed 98.0–99.0% identity, 3 showed 97.0–98.0% identity, 1 showed 96.0–97.0% identity, and 1 showed 95.0–96.0% identity.

The f-element was verified through alignment of Sanger sequences with the expected f-element nuclear insert microsatellite region [5].

### Data Analysis and Visualization

Data visualization for SRD distribution, haplotype frequency, and symbiont prevalence across samples was performed using DataClassroom, with data organized in Microsoft Excel. Geographic mapping of sample locations was generated using DataWrapper. Statistical analyses, including Fisher’s Exact tests, were conducted using GraphPad Prism.

## Data Availability

Representative sequences were selected to capture each observed COI haplotype, *Wolbachia* strain, and f-element detection, with additional geographic representation included for common haplotypes. All data supporting the findings of this study, including sequence data and associated metadata, are publicly available on Zenodo at https://doi.org/10.5281/zenodo.19867604

## Results

We examined 295 *A. vulgare* individuals (180 females and 115 males) from 12 U.S. states across a broad geographic range (Figure 1) for the presence of Wolbachia and the f-element (Table 1). Overall, 43% of individuals carried at least one sex ratio distorter (SRD). Among all individuals, 24% were infected with *w*VulC (*n* = 72) and 5% with *w*VulM (*n* = 15). The f-element was detected in 14% of individuals. SRD presence differed between sexes, as *Wolbachia* was detected in both males and females, whereas the f-element was detected only in females.

**Figure 1:**
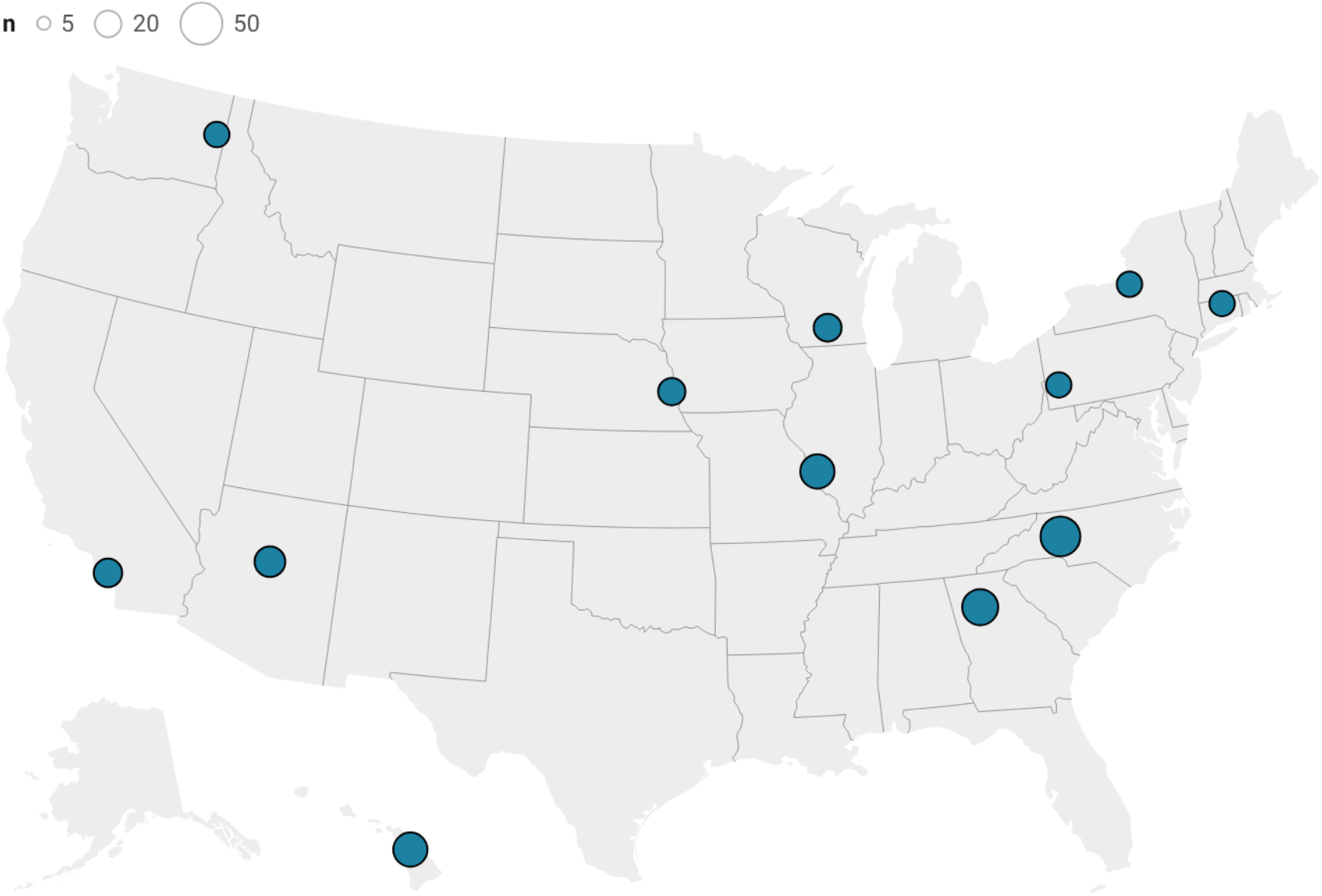
*Armadillidium vulgare* sampling locations across the United States, highlighting the geographic extent of sampling. Circle size is proportional to the number of individuals sampled at each location. Most states were represented by a single sampling location, while California, Georgia and New York each contained two sites (see Table 1 for complete data).

SRD distribution of *Wolbachia* and the f-element varied across the sampled states (Figure 2). Several states showed relatively high frequencies of *Wolbachia*, including North Carolina (55.6%), New York (75%), Connecticut (68.8%), Washington (43.8%), and Wisconsin (40%). In contrast, Missouri and Nebraska showed higher relative contributions of the f-element compared to *Wolbachia* (Missouri: 31.3% vs. 3.13%; Nebraska: 26.3% vs. 5.26%). Other states, including Georgia (f-element 22.2%, *Wolbachia* 27.8%), California (28.6%, 23.8%), and Pennsylvania (25%, 31.3%), showed mixed f-element/*Wolbachia* profiles, with both detected at similar frequencies. Arizona had less than 5% SRD presence, and Hawaii was the only state sampled in which no SRDs were detected. *Wolbachia* and the f-element co-occurred in 9 out of 12 states sampled.

**Figure 2:**
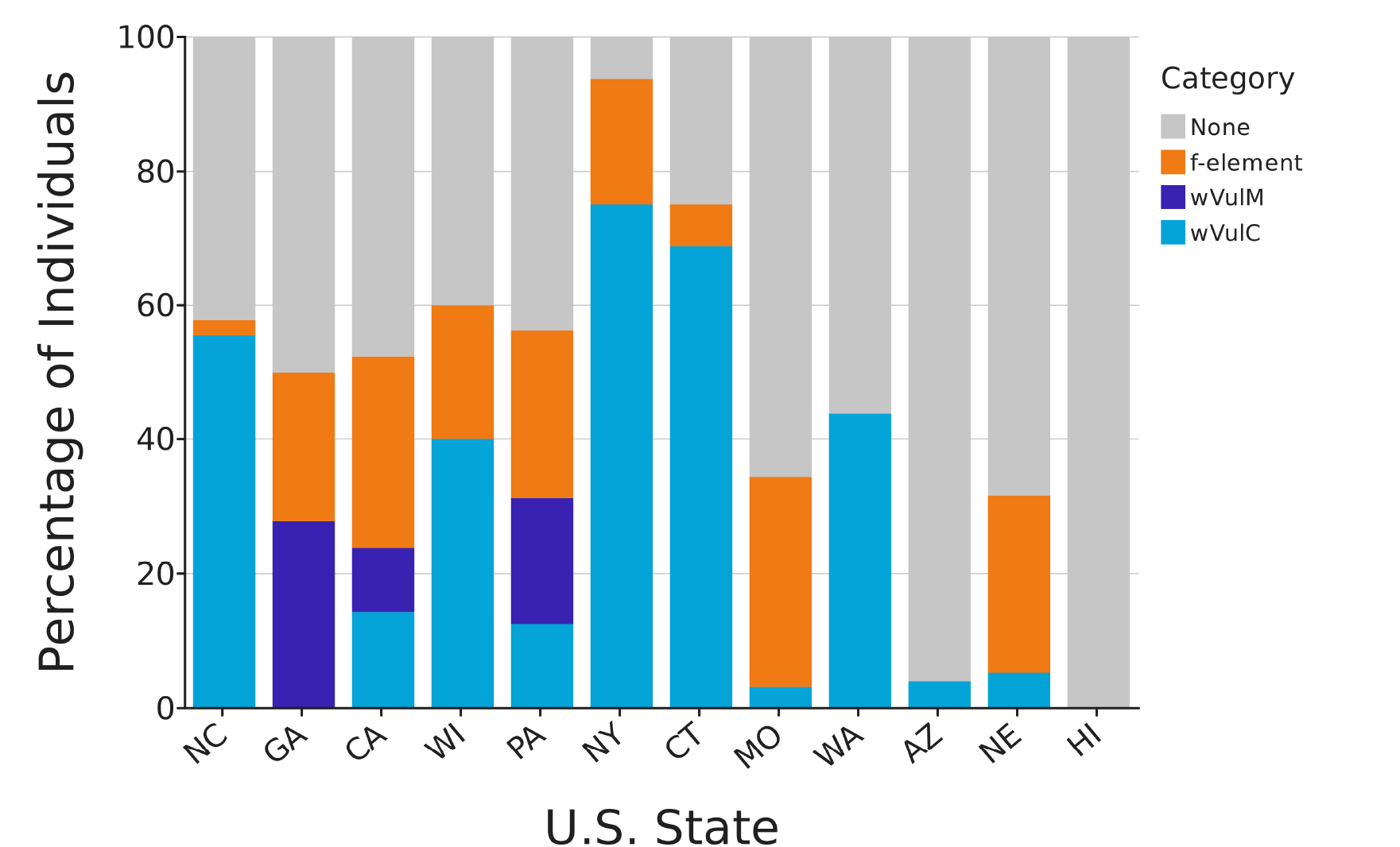
Distribution of sex ratio distorters (SRDs) across U.S. states. Stacked bars show the proportion of individuals carrying *Wolbachia* strains (*w*VulC and *w*VulM), the f-element, or no detectable SRD. Each bar is normalized to 100% of individuals within that population.

COI sequencing revealed six haplotypes across the sampled states (I, II, IV, V, VI, and XV) (Table 2). Two haplotypes were most common overall, with haplotype V accounting for 42.3% of individuals and haplotype VI for 47.7%. The remaining haplotypes were less frequent, including II (7.1%), I (1.3%), IV (1.0%), and XV (0.3%).

Haplotype V was most abundant in North Carolina (93%), New York (75%), and Connecticut (87.5%), whereas haplotype VI was most abundant in California (76.2%), Missouri (68.8%), Arizona (84%), Nebraska (94.7%), and Hawaii (57.6%). In all states sampled, haplotypes V and VI co-occurred. In Georgia and Pennsylvania, haplotype II was also relatively common, comprising 41.7% and 31.3% of individuals, respectively. Haplotypes I, IV, and XV were rare across all sampled populations.

Several states, including North Carolina, Georgia, California, Wisconsin, Pennsylvania, and Missouri, exhibited three or more haplotypes, indicating greater haplotype diversity. In contrast, New York, Connecticut, Washington, Arizona, Nebraska, and Hawaii were composed primarily of haplotypes V and VI (Figure 3).

**Figure 3.**
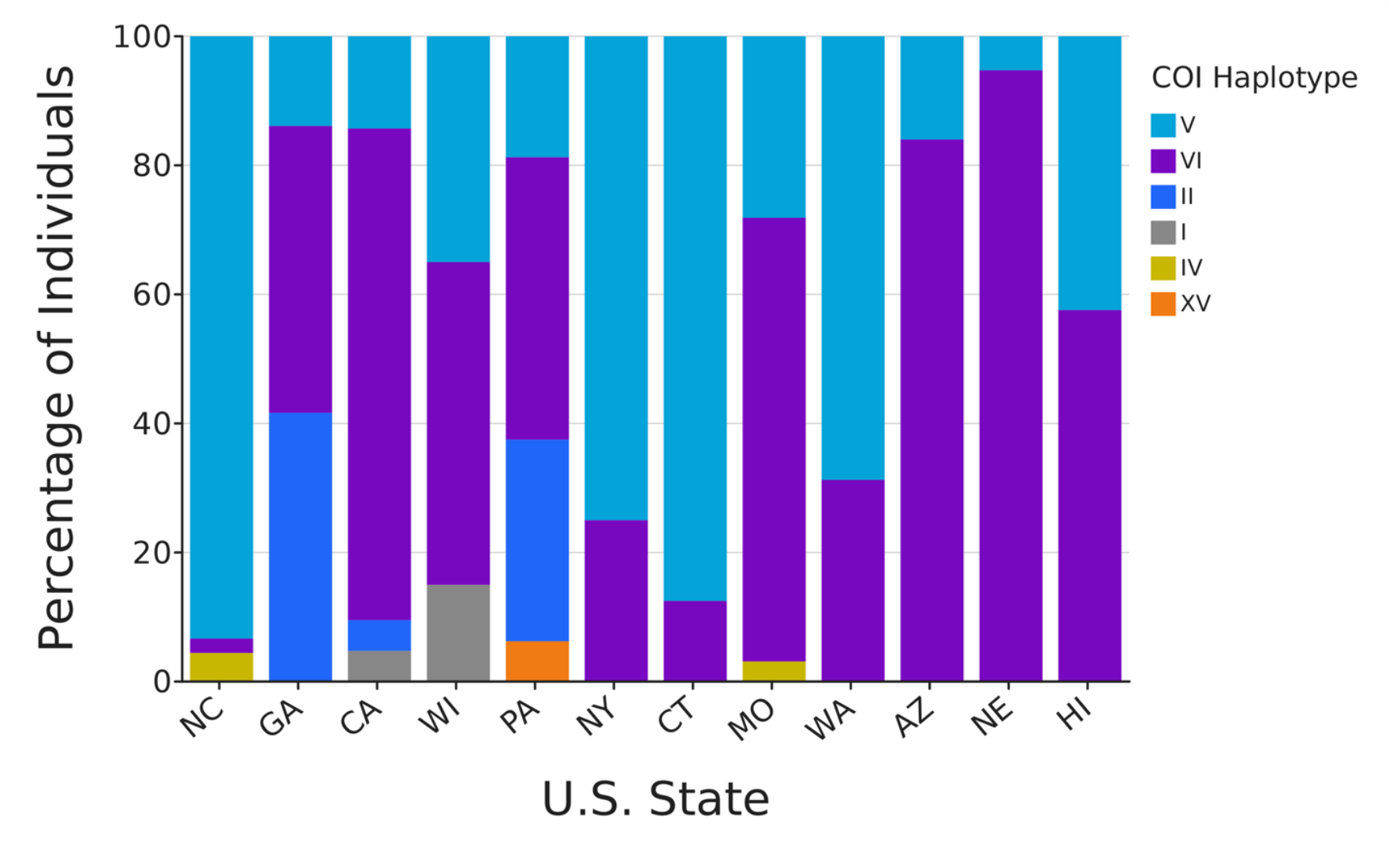
Distribution of COI haplotypes across U.S. states. Stacked bars show the percentage of individuals within each state assigned to each COI haplotype (V, VI, II, I, IV, XV). Each bar is normalized to 100% of individuals per state. Haplotype composition varied among states, with haplotypes V and VI predominating in most populations, while some states exhibited greater haplotype diversity (≥3 haplotypes). Sample sizes per state are provided in Table 1.

Comparison of COI haplotype and SRD occurrence revealed several clear and consistent patterns (Figure 4). Haplotypes V and VI contained all three SRD types (*w*VulC, *w*VulM, and f-element). Haplotype II was primarily associated with *Wolbachia* strain *w*VulM, while haplotypes I and IV were more frequently associated with wVulC. The f-element was detected in 27.7% of individuals in haplotype VI and less commonly in haplotypes II (9.5%) and V (0.8%) (Figure 4).

**Figure 4:**
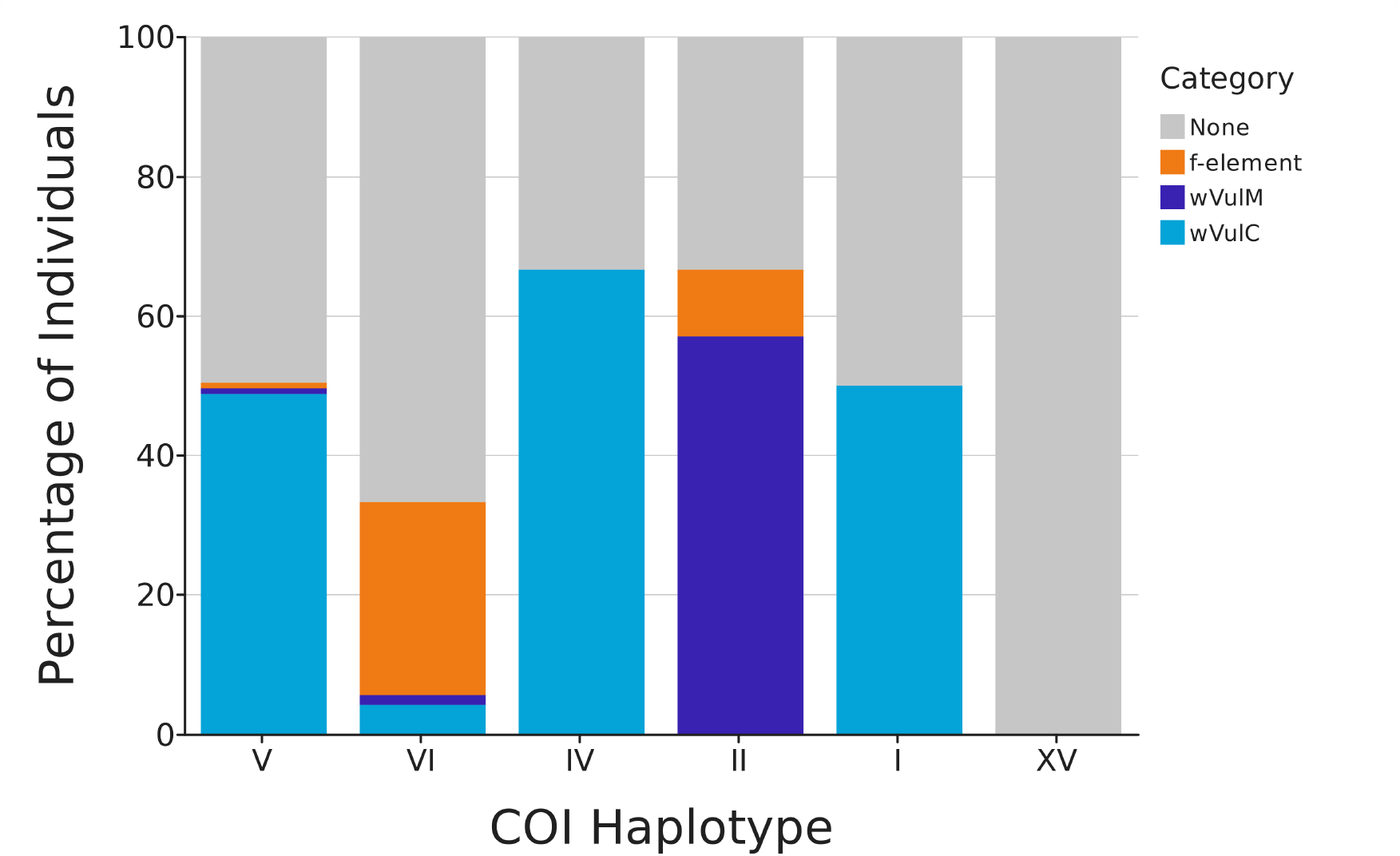
Distribution of sex ratio distorters (SRDs) according to COI haplotype. Stacked bars show the proportion of individuals carrying *Wolbachia* strains *(w*VulC and *w*VulM), the f-element, or no detectable SRD. Each bar is normalized to 100% of individuals within each haplotype. Sample sizes: V (n = 125), VI (n = 141), IV (n = 3), II (n = 21), I (n = 4), and XV (n = 1) (total n = 295). The f-element is primarily associated with haplotype VI, while *Wolbachia* shows broader distribution across haplotypes.

A Fisher’s exact test was used to compare the frequency of the f-element between haplotype VI and all other haplotypes. The null hypothesis was that the f-element frequency is independent of haplotype, and no difference in frequency would be expected between haplotype VI and other haplotypes. The frequency of the f-element in haplotype VI was significantly higher than all other haplotypes combined (Fisher’s exact test, p<0.0001).

## Discussion

Sampled U.S. populations exhibit highly variable SRD distribution patterns, which are consistent with multiple, independent introductions, rather than a single spreading lineage. *Wolbachia* distribution is heterogeneous across states (Figure 1). Strain *w*VulM is primarily associated with COI haplotype II, and strain *w*VulC is distributed across multiple haplotypes (V, VI, and I). Because both *Wolbachia* and mitochondrial DNA are maternally inherited, they are expected to associate with specific COI haplotypes. In a French population, *Wolbachia* has been shown to influence mitochondrial genetic structure through hitchhiking, producing linkages between *Wolbachia* strains and mitochondrial haplotypes [14]. In this way, maternally inherited symbionts can change the distribution of associated mitochondrial haplotypes and reduce mitochondrial diversity, strengthening the association between *Wolbachia* and mitochondrial haplotypes [15, 16, 17]. However, this trend is not consistently observed in U.S. populations, suggesting that *Wolbachia* distribution may be influenced by the mixing of multiple mitochondrial lineages and variation in *Wolbachia* strain composition within populations [18], as shown in Figure 2.

This inconsistency is expected in introduced systems where multiple introductions and population mixing can disrupt expected genetic associations, producing heterogeneous patterns [19,20]. European populations, representing the ancestral range of *A. vulgare*, show stronger associations between *Wolbachia* and COI [9], consistent with a more stable, near-equilibrium system. Weaker associations seen in the sampled U.S. population may reflect more recent introduction history and ongoing population mixing.

The evolutionary origin and persistence of the f-element in *A. vulgare* reflect a complex interaction between host genetics and the horizontal transfer of symbiont DNA [6]. In contrast to the maternally inherited *Wolbachia*, the f-element is a nuclear element and is not expected to be strictly associated with mitochondrial haplotypes. However, in the U.S., the f-element shows a strong and statistically significant association with haplotype VI (Figure 4). Although haplotype VI represents approximately one-third of individuals in U.S. populations, the f-element was largely restricted to this haplotype. This relationship is supported by the Fisher’s Exact test (p<0.0001) and indicates a non-random association between the f-element and mitochondrial background. In most populations with the f-element, multiple mitochondrial haplotypes were observed, indicating that these samples were genetically diverse. Despite this genetic diversity, the f-element remained largely restricted to haplotype VI across populations, suggesting that the observed association reflects a population-level pattern rather than a simple sampling artifact.

One possibility for this pattern is that individuals carrying the f-element were introduced from European source populations in which haplotype VI was already present, leading to an initial association between nuclear and mitochondrial elements. In contrast, European populations do not show a similar association between the f-element and haplotype VI [9], suggesting that introduction history likely contributed to the initial association. However, even though haplotype VI is common, that alone does not explain why the f-element is so strongly found in that haplotype. It is possible that processes within U.S. populations may help maintain this association. For example, genetic drift in small, structured populations may increase the frequency of the f-element associated with specific haplotypes.

In U.S. populations, *Wolbachia* shows variable and mixed distributions across haplotypes, while the f-element remains strongly associated with haplotype VI, indicating that these SRDs are being shaped by different processes.

## Limitations

This study has several limitations that should be considered when interpreting the results. Crowdsourced sampling across U.S. states was uneven, and sample sizes within individual locations were often small, which may influence estimates of haplotype and SRD frequencies. Although multiple haplotypes were observed in most populations, some degree of local relatedness cannot be excluded. Additionally, comparisons to European populations are based on previously published studies and not from direct analysis of their datasets. This limits our ability to quantitatively assess differences between native and introduced ranges. While a strong association between the f-element and haplotype VI was observed, we cannot determine whether this pattern reflects introduction history or biological processes within U.S. populations. Because sampling was not designed to estimate population-level sex ratios and instead emphasized representation of SRD diversity across locations, we did not formally assess relationships between SRD prevalence and sex ratio across states.

## Acknowledgements

We thank those who contributed to our crowdfunding campaign through Experiment.com, Scott Kent (Principal of FCS Innovation Academy STEM High School), Dr. Richard Cordaux, Harshita Nekkanti, and Varshini Ganesh Mohan. We are grateful to the individuals who contributed pillbug samples to this study: Jessica Diaz (PA), Parks Collins (NC), Subodh Niroula (CA), Shaun Cross (NE), Liz Cowles (CT), Deborah Beam (NY), Chris Chandler (NY), Sarah Holmes (MO), and Dan Shay (WA). We also thank the Bordenstein Lab and the *Wolbachia* Project for PCR primer support.

